## Supplemental Info for "Mapping Serotonergic Dynamics using Drug-Modulated Molecular Connectivity"

### Supplementary Methods

#### Schiffer brain atlas

The Schiffer brain atlas (implemented in PMOD data analysis software) was used to parcelate the brains for ROI seed-based analysis into 23bilateral regions, the periaqueaductal gray (PAG) and the septum (Sep), resulting in a total number of 48 brain regions. A list of all regions and their respective abbreviations is provided in Supplementary Table 1.

Supplementary Table 1: Brain regions included in the Schiffer rat brain atlas, including their respective volumes and abbreviations.

| **Brain region (ROI)** | **Hemisphere** | **ROI volume [mm^3^]** | **Position on correlation matrix** | **Abbreviation** |
| --- | --- | --- | --- | --- |
| Nucleus Accumbens | left | 7.944 | 1 | NAc |
|  | right |  | 2 |  |
| Amygdala | left | 21.120 | 3 | Amyg |
|  | right |  | 4 |  |
| Dorsal Striatum | left | 43.552 | 5 | Str |
|  | right |  | 6 |  |
| Auditory Cortex | left | 27.520 | 7 | Au |
|  | right |  | 8 |  |
| Cingulate Cortex | left | 14.480 | 9 | Cg |
|  | right |  | 10 |  |
| Entorhinal Cortex | left | 59.016 | 11 | Ent |
|  | right |  | 12 |  |
| Insular Cortex | left | 21.128 | 13 | Ins |
|  | right |  | 14 |  |
| Medial Prefrontal Cortex | left | 6.304 | 15 | mPFC |
|  | right |  | 16 |  |
| Motor Cortex | left | 32.608 | 17 | M1 |
|  | right |  | 18 |  |
| Orbitofrontal Cortex | left | 18.936 | 19 | OFC |
|  | right |  | 20 |  |
| Parietal Cortex | left | 7.632 | 21 | PaC |
|  | right |  | 22 |  |
| Retrosplenial Cortex | left | 18.920 | 23 | RS |
|  | right |  | 24 |  |
| Somatosensory Cortex | left | 71.600 | 25 | S1 |
|  | right |  | 26 |  |
| Visual Cortex | left | 36.136 | 27 | V1 |
|  | right |  | 28 |  |
| Anterodorsal Hippocampus | left | 25.064 | 29 | CA1 |
|  | right |  | 30 |  |
| Posterior Hippocampus | left | 9.784 | 31 | CA1-p |
|  | right |  | 32 |  |
| Hypothalamus | left | 18.352 | 33 | Hyp |
|  | right |  | 34 |  |
| Olfactory Cortex | left | 14.008 | 35 | OC |
|  | right |  | 36 |  |
| Superior Colliculus | left | 7.136 | 37 | SC |
|  | right |  | 38 |  |
| Midbrain | left | 11.448 | 39 | MB |
|  | right |  | 40 |  |
| Ventral Tegmental Area | left | 5.528 | 41 | VTA |
|  | right |  | 42 |  |
| Inferior Colliculus | left | 5.744 | 43 | IC |
|  | right |  | 44 |  |
| Thalamus | left | 30.712 | 45 | Th |
|  | right |  | 47 |  |
| Periaqueaductal Gray | - | 9.904 | 47 | PAG |
| Septum | - | 9.36 | 48 | Sep |

#### Radiotracer synthesis

[^11^C]DASB was synthesized similar to the procedure reported by Wilson et al. (Wilson et al., 2000). Briefly, [^11^C]MeI was trapped in a solution of 2 mg precursor in 500 µl DMSO. After heating to 100 °C for 2 min the reaction was diluted with 1.5 ml HPLC eluent (3 mM Na_2_HPO_4_ containing 64 % MeCN) and purified on a Luna C18(2) column (250 mm x 10 mm, Phenomenex). The isolated peak was diluted with 70 ml water containing 20 mg sodium ascorbate, loaded onto a conditioned Strata-X cartridge (Phenomenex), eluted with 0.5 ml ethanol and diluted with 5 ml phosphate-buffered saline.

#### DVR-1 detrending

We applied a piecewise linear detrend to remove remaining tracer uptake dynamics from the dynamic DVR-1 time-series and its effect on the artificial inflation of correlation values in the early parts of the scan, before reaching a steady-state, while only discarding the first 20 minutes of every scan, which were too strongly dominated by tracer uptake effects (Supplementary Figure 1).

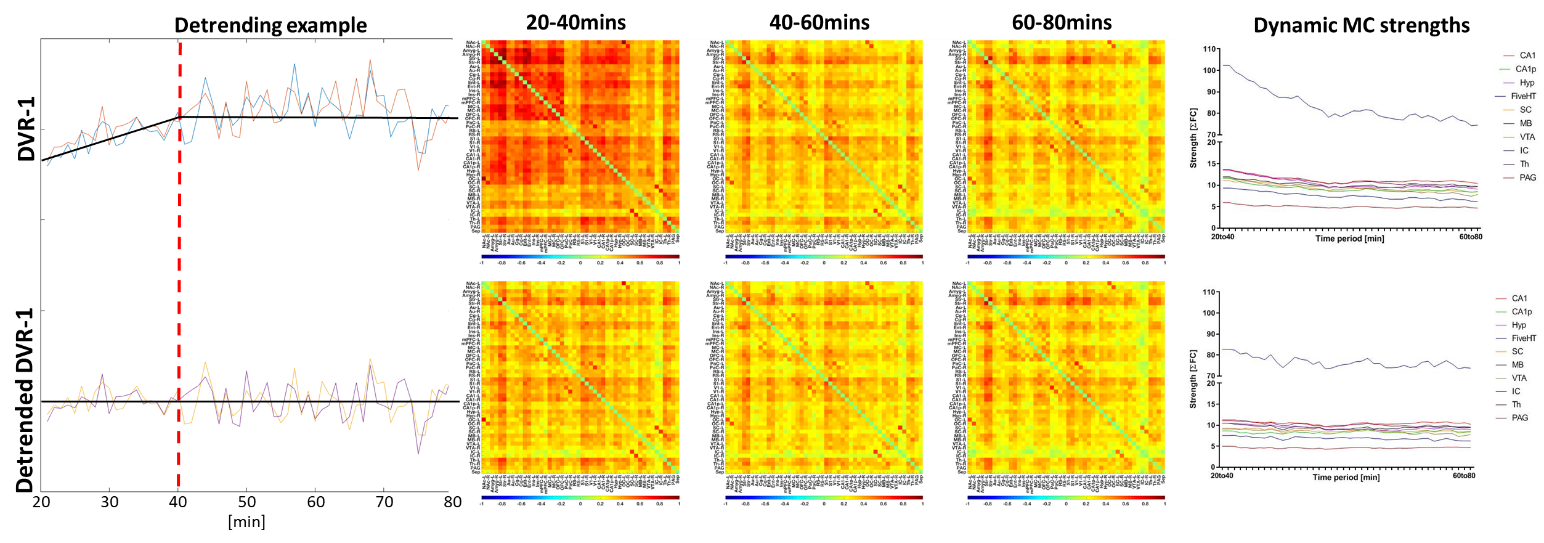

Supplementary Figure 1: Detrending procedure of DVR-1 time-courses. Left panel: exemplary dynamic DVR-1 timecourses between minute 20 and 80 after scan start. An increasing trend can be observed until minute 40 for the raw DVR-1 timecourses. After detrending the segments 20-40 and 40-80 separately, a linear time-series can be obtained. Middle panel: group-level MC correlation matrices before and after detrending of the DVR-1 time-series. Without detrending, MC values between minutes 20 and 40 are strongly inflated, this aspect being solved by the applied detrending procedure. Right panel: without detrending, the MC strengths only reach temporally constant values towards the end of the scans, therefore not allowing the detection of a potential intervention prior to that timepoint. When applying detrending, the strengths remain already from the time period 20-40mins onward.

Schiffer, W.K., Mirrione, M.M., Biegon, A., Alexoff, D.L., Patel, V., Dewey, S.L., 2006. Serial microPET measures of the metabolic reaction to a microdialysis probe implant. J Neurosci Methods 155, 272-284.

Wilson, A.A., Ginovart, N., Schmidt, M., Meyer, J.H., Threlkeld, P.G., Houle, S., 2000. Novel Radiotracers for Imaging the Serotonin Transporter by Positron Emission Tomography:  Synthesis, Radiosynthesis, and in Vitro and ex Vivo Evaluation of 11C-Labeled 2-(Phenylthio)araalkylamines. Journal of Medicinal Chemistry 43, 3103-3110.
